# Radiation dose effects in correlative X-ray / cryo-electron microscopy of frozen hydrated biological samples

**DOI:** 10.1101/2025.10.07.680863

**Authors:** Thorsten B. Blum, Vincent Olieric, Ana Diaz, Takashi Ishikawa, Volodymyr M. Korkhov

**Affiliations:** PSI Center for Life Sciences, Paul Scherrer Institute, 5232 Villigen PSI, Switzerland; PSI Center for Photon Science, Paul Scherrer Institute, 5232 Villigen PSI, Switzerland; Department of Biology, ETH Zurich, Zurich 8093, Switzerland; Institute of Molecular Biology and Biophysics, ETH Zürich, Otto-Stern-Weg 5, Zürich 8093, Switzerland

**Keywords:** cryo-X-ray tomography, cryo-electron microscopy, correlative microscopy

## Abstract

In cryo-electron microscopy (cryo-EM), imaging of biological specimens is restricted by the limited field of view and by sample thickness. Hard X-ray imaging, with its ability to penetrate samples several tens of micrometers thick, offers a complementary approach for high-resolution visualization. A major concern is whether cryo-preserved samples can withstand the handling conditions at synchrotron facilities without excessive icing, and whether the radiation exposure during X-ray imaging compromises specimen integrity, thereby hindering subsequent attempts to achieve high-resolution 3D reconstructions via cryo-EM. To evaluate this, we deposited apoferritin samples on a cryo-EM grid, exposed them to varied X-ray doses typical for X-ray tomography experiments at a synchrotron facility, and subsequently analysed the exposed particles by cryo-EM. Despite the apparent damage sustained throughout the experiment, the samples remained amenable to cryo-EM analysis, with structural details at a resolution of ∼4 Å at the highest absorbed X-ray dose of 100 MGy. By comparison, a similar cryo-EM dataset of the apoferritin particles that were not exposed to X-rays but were mounted on the same cryo-EM grid, resulted in a 3D reconstruction at 3.17 Å resolution. Thus, while radiation damage may limit the high-resolution information in specimens processed by cryo-X-ray tomography, the cryo-preserved biological material exposed to these high X-ray doses can be still used for subsequent cryo-EM workflows aiming to obtain structural biology insights at intermediate to high resolution. These findings lay the groundwork for an integrated imaging workflow that combines X-ray and cryo-EM techniques to enable multiscale analysis of thick vitrified biological specimens.

## Introduction

Cryo-electron microscopy (cryo-EM) is a foundational technique in structural biology, renowned for its ability to produce high-resolution data of biological specimens (Kühlbrandt, 2014; Saibil, 2022). It allows researchers to visualize intricate cellular architectures and protein complexes, thereby deepening our insight into complex biological mechanisms. However, achieving multiscale high-resolution imaging with cryo-EM remains challenging due to constraints such as a limited field of view and the necessity for very thin samples amenable to transmission EM analysis.

Given the limited field of view of cryo-EM, employing multiscale imaging techniques before cryo-EM is essential for effective sample characterization and for targeting the regions of interest. This process remains challenging, not only because of the restricted imaging area but also due to the need for thin, electron-transparent frozen-hydrated specimens. To overcome these hurdles, correlative light and electron microscopy (CLEM) has proven to be a valuable approach for pinpointing regions of interest within frozen biological samples, whether at room temperature (Shafiei *et al*., 2025) or under cryogenic conditions (Klein *et al*., 2021).

However, despite their utility, CLEM approaches rely on fluorescence of the objects of the target structures to identify the region of interest when employing the waffle method, in which the sample is vitrified on an EM grid through high-pressure freezing (Kelley *et al*., 2022; Klykov *et al*., 2022). X-ray imaging presents a compelling alternative, offering the capacity to image thick, unstained samples across a broader field of view. The integration of hard X-ray nano-tomography (Shahmoradian *et al*., 2017; Longo *et al*., 2020; Kuan *et al*., 2020) with cryo-EM may facilitate the visualization of large and complex biological specimens with volumes up to 100 × 100 × 100 µm^3^, while also allowing for the precise targeting of specific regions for detailed analysis using cryo-electron tomography (cryo-ET).

To integrate the various techniques, we aim to develop a workflow of processes to achieve this in the long term (Figure 1). For this purpose, the biological samples must first undergo cryo-fixation under cryogenic conditions, which can consist of plunge freezing in liquid ethane, as is done for samples thinner than about 10 𝜇m and which can be imaged by cryo soft X-ray tomography (Groen *et al*., 2019; Guo & Larabell, 2019; Groen *et al*., 2025). However, for samples thicker than about 10 𝜇m, the sample preparation may require high-pressure freezing followed by trimming with an ultramicrotome (Figure 1A) (Holler *et al*., 2017). Hard X-ray nano-tomography can then be employed to locate regions of interest (Figure 1B), which may be thinned using focused ion beam (FIB) (Marko *et al*., 2006; Rigort *et al*., 2012) milling or cryo-electron microscopy of vitreous sections (CEMOVIS) (Studer *et al*., 2001; Al-Amoudi *et al*., 2004). Finally, these thinned areas can be imaged using cryo-EM, aiming to obtain 3D reconstructions with high (sub-nanometer) spatial resolution (Figure 1C). Figure 1 shows the envisioned workflow for correlative hard X-ray nano-tomography of tissues and cryo-EM.

**Figure 1.**
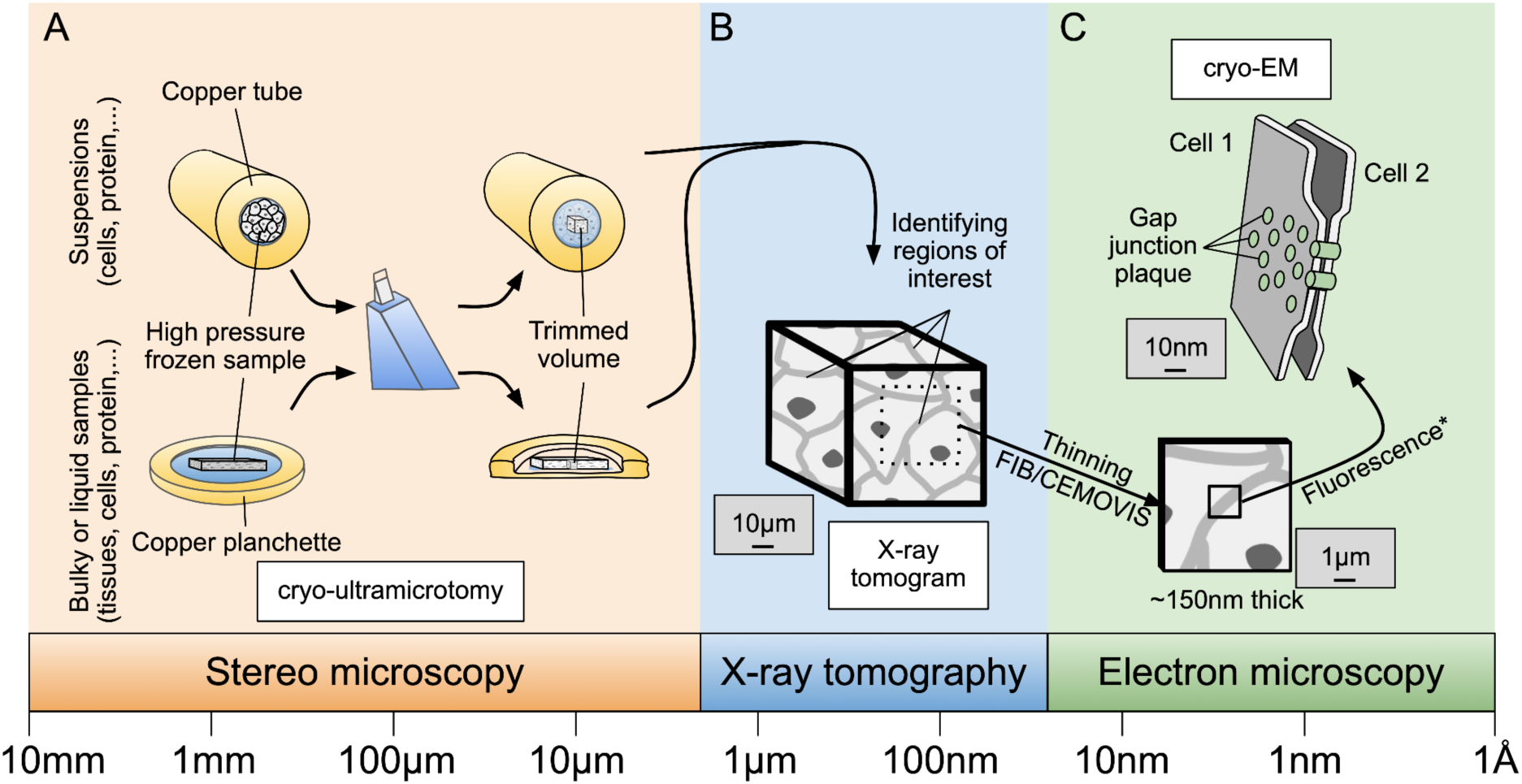
Schematic workflow of the process envisioned for correlative hard X-ray nano-tomography and cryo-EM. The purpose of this workflow is to identify a region of interest (e.g., a gap junction plaque at the cell-cell interface, illustrated here). To achieve this, (A) cells are subjected to high-pressure freezing within copper tubes (inner diameter: 350 µm), or tissues are embedded in copper planchettes with a 200 µm cavity. These samples are then precisely trimmed using a diamond knife to yield a rectangular section measuring approximately 70 × 70 µm^2^ in area and 30 µm in thickness, optimized for hard X-ray nano-tomography. (B) An X-ray tomogram of the trimmed volume is acquired, allowing identification of regions of interest (ROIs). (C) Thin sample layers are then extracted from selected ROIs using either cryo-electron microscopy of vitreous sections (CEMOVIS) or focused ion beam (FIB) milling (*optionally guided by fluorescence light microscopy if fluorescence markers are present) for downstream cryo-EM analysis.

To the best of our knowledge, correlative imaging combining cryo hard X-ray nano-tomography and cryo-ET has not been successfully performed to date. Only individual steps of the entire pipeline (Figure 1) under room temperature or cryo conditions have been demonstrated to confirm their feasibility. For example, it has been shown that cryo-preserved samples can be prepared and successfully analysed by X-ray nano-tomography (Holler *et al*., 2017). Although the tomograms of frozen samples were not correlated for region of interest (ROI) extraction, studies involving fixed biological samples have demonstrated that ROIs identified through micro-computed tomography (micro-CT) can be trimmed with a microtome and subsequently transferred to a focused ion beam scanning electron microscope for imaging in scanning electron microscopy (SEM) mode (Karreman *et al*., 2016).

Other studies have demonstrated that cryo-light/fluorescence can be correlated with cryo soft X-ray tomography (Okolo *et al*., 2021). Likewise, light/fluorescence correlated with hard X-ray tomography (correlative single-cell hard X-ray computed tomography and X-ray fluorescence imaging) or correlating soft X-ray tomography under non-cryo conditions with non-biological specimens using STEM has been demonstrated (Lo *et al*., 2019). While these developments mark significant progress, they do not address key questions surrounding the integration of cryo-X-ray nano-tomography with cryo-ET. Crucially, uncertainties remain about: (i) whether cryo-preserved samples can endure the handling at the synchrotron facilities without becoming excessively icy; (ii) whether the radiation doses absorbed during X-ray imaging compromise the specimen’s integrity, limiting any subsequent efforts aimed at obtaining high-resolution 3D reconstructions by cryo-EM.

X-ray damage is classified into primary and secondary damage (Henderson, 1995; Nass, 2019). Primary damage results from the direct interaction between the radiation beam and electrons, resulting in the ejection of energetic electrons from atoms through processes such as the photoelectric effect, Auger effect, and Compton scattering. These events can break chemical bonds, causing molecular degradation. Secondary damage arises from the reactions of radiolytic products, such as free radicals generated by these energetic electrons. This radiation-induced damage results in the degradation of image resolution during subsequent EM processing.

Radiation damage has been previously extensively investigated using X-ray crystallography of crystalline samples (Garman & Weik, 2023). Early insights were provided by Henderson (1990) (Henderson, 1990), who predicted that exposure to X-ray doses of approximately 20 MGy at nitrogen temperatures would destroy half of the intensity of crystalline diffraction patterns in protein crystals, supported experimentally by Gonzalez & Nave (1994) (Gonzalez & Nave, 1994). Furthermore, Teng and Moffat (2000) demonstrated through diffraction experiments that lysozyme crystals could withstand up to 16 MGy and still achieve a resolution of 1.6 Å (Teng & Moffat, 2000). Additionally, Owen et al. (2006) reported that apoferritin crystals exhibited significant intensity loss – 50% or more – when subjected to 43 MGy for a resolution range of 2.6–2.5 Å, or 200 MGy for lower-resolution ranges of 20–10 Å (Owen *et al*., 2006). In single proteins, high-resolution structural information is particularly susceptible to damage, as radical formation from X-ray exposure often compromises sensitive side chains (Teng & Moffat, 2000; Owen *et al*., 2006). How the X-ray dose affects the amorphous frozen samples that more closely resemble the typical biological samples (cells, tissues, etc.) has not been evaluated so far, to the best of our knowledge.

Here we tested whether exposure of biological samples to radiation doses typically used in hard X-ray nano-tomography (Shahmoradian *et al*., 2017) would still preserve the structural information at high resolution – a prerequisite for developing a cryo-correlative workflow for combining X-ray imaging with cryo-EM. Typical doses reported in the literature for frozen hydrated tissues are between 1 and 20 MGy (Shahmoradian *et al*., 2017; Holler *et al*., 2017), however, we expect that doses up to 100 MGy could be absorbed, e.g. by ptychographic X-ray computed tomography at 4th-generation synchrotron sources. An exposure of 100 MGy required for recording a ptychographic X-ray computed tomogram corresponds to a pre-exposure of approximately 27 e^-^/Å^2^ when using a 300 keV electron microscope with a conversion factor of 3.7 (Dickerson *et al*., 2024; Groen *et al*., 2025). Cryo-EM studies have demonstrated that datasets exposed to ∼19 e^-^/Å^2^ or ∼24 e^-^/Å^2^ at 300 keV, after discarding the initial movie frames, still yielded reconstructions at 3.35 Å (Grant & Grigorieff, 2015) and 3.94 Å (Allegretti *et al*., 2014), using particle alignments derived from the full, dose-weighted movies. Experiments assessing pre-exposure with X-rays or electrons suggest that, following the acquisition of a ptychographic X-ray computed tomogram, the resolution should remain sufficient to trace the protein backbone.

However, in cryo-correlative workflows that combine X-ray imaging with cryo-EM, it is essential to consider the cumulative radiation dose absorbed by biological specimens during both modalities. To assess the radiation effects, we chose an *in vitro* sample, apoferritin, as a particularly suitable specimen that can be frozen on cryo-EM grids in thin ice, yields high-resolution density maps in cryo-EM analysis and should thus enable precise assessment of the combined effects of X-rays and electrons on biological structures.

## Methods

### Apoferritin on EM grids

Purified apoferritin (30 µl) derived from equine spleen (Sigma-Aldrich) was desalted using a micro bio-spin column (Bio-Rad) into a Tris buffer (20 mM Tris/HCl pH 7.5, 300 mM NaCl). Aliquots of 3 µl of apoferritin at a concentration of ∼0.85 mg/ml were applied to glow discharged Quantifoil Cu200 R1.2/1.3 holey carbon EM grids or Quantifoil Au200 R2/2 holey carbon EM finder grids. Grids were blotted for 2 seconds with a blot force of 10 and plunge-frozen in liquid ethane using a Vitrobot (Thermo Fisher Scientific) with the environmental chamber set at 100% humidity and 8 °C. Grids prepared in this way were stored in liquid nitrogen before imaging.

### X-ray radiation exposure

EM grids were mounted in AutoGrid rings and secured with C-clips (Thermo Fisher Scientific). These clipped grids were then placed into holders compatible with SPINE vials (Huang *et al*., 2020) and shipped to the European Synchrotron Radiation Facility (ESRF) in Grenoble, France. Using a vacuum cryo transfer system in liquid nitrogen (LN₂), the sample holders were transferred to the loading carrier. Exposures were conducted at 100 K at the ID30B beamline of the ESRF, using a photon energy of 13.5 keV, a beam size of 30 × 30 µm^2^, and a flux of 1.0 × 10^13^ ph/sec. Areas of 90 × 90 µm^2^ field of view were exposed using the mesh scan function of MXCuBE (Bowler *et al*., 2010). The beam follows a continuous snake-like trajectory, resulting in estimated radiation doses of 1 MGy and 100 MGy, while the maximum variation in the exposed volume was around 9 %. The radiation dose was calculated using Formula 5 from Holton’s “A beginner’s guide to radiation damage” (Holton, 2009), without accounting for the sample thickness, and applying exposure times of 0.213 sec and 21.3 sec.

### Cryo-TEM imaging

Data were acquired on a Titan Krios G3i electron microscope at 300 keV (Thermo Fisher Scientific), with a GIF BioQuantum filter (20 eV slit width) and a K3 Summit electron counting direct detection camera (Gatan) at a magnification of 76,923 ×, resulting in a pixel size of 0.65 Å, using EPU (Thermo Fisher Scientific). The defocus varied between –0.6 and –2.4 µm. Overall, 1,730 movies were recorded with a total dose of 60 e^-^/Å^2^ per movie (0.92 s exposure in total, 40 frames in total). Of these, for 0, 1, and 100 MGy, the number of movies was 760, 405, and 565, respectively (Table 1).

**Table 1.** Cryo-EM structure determination.

| **Data collection** | **0 MGy** | **1 MGy** | **100 MGy** |
| --- | --- | --- | --- |
| Magnification | 76,923 x |  |  |
| Pixel size (Å) | 0.65 |  |  |
| Defocus Range (µm) | -0.6 to -2.4 |  |  |
| Voltage (kV) | 300 |  |  |
| Exposure time (s) | 0.92 |  |  |
| Total dose (e/Å<sup>2</sup>) | 60 |  |  |
| Number of frames (tiff) | 40 |  |  |
| Dose per frame (e/Å<sup>2</sup>) | 1.5 |  |  |
| Number of movies | 760 | 405 | 565 |

| **Reconstitution** | **0 MGy** | **1 MGy** | **100 MGy** |
| --- | --- | --- | --- |
| Automatically picked particles | 60,105 | 48,194 | 80,664 |
| After 2D classification (particles) | 33,386 | 8,548 | 13,728 |
| After 3D classification (particles) | 30,265 | 6,488 | 9,547 |
| Final overall resolution (Å) | 2.83 | 3.51 | 3.71 |
| Estimated map sharpening B-factor (Å<sup>2</sup>) | -80 | -99 | -100 |
| Randomly equally picked particles | 6,488 | 6,488 | 6,488 |
| EMDB accession number | 55343 | 55338 | 55344 |
| Final overall resolution (Å) | 3.17 | 3.56 | 3.88 |
| Estimated map sharpening B-factor (Å<sup>2</sup>) | -63 | -83 | -97 |

### Image processing / Data analysis

For image processing, the single particle cryo-EM analysis software package Relion 5 (Scheres, 2012) was used. The movies were drift-corrected and dose-weighted using RELION’s own implementation of the MotionCor2 (Zheng *et al*., 2017) algorithm, and the contrast transfer function parameters were estimated from the motion-corrected electron micrographs using CTFFIND4 (Rohou & Grigorieff, 2015). Apoferritin particles were automatically picked by using three 2D classes of apoferritin. Boxes with a size of 512 pixels, rescaled to 64 pixels, were extracted. Several rounds of 2D classification and one 3D classification were done. For the first 3D classification a map obtained from EMPIAR 11273 was used as initial reference. The reference was low-pass filtered to 60 Å and an octahedral symmetry was applied for all 3D runs. Only good classes were extracted with boxes of 400 pixels, without re-scaling. An additional round of 3D classification was performed, followed by 3D refinement and post-processing using automatic B-factor estimation. For post-processing, a mask was generated in Relion 5 from the refinement maps by extending the binary maps by 10 pixels and adding a soft-edge by 20 pixels. Subsequently, Bayesian polishing and CTF refinement were carried out to estimate (i) higher-order aberrations, including beam tilt, trefoil, and fourth-order aberrations; (ii) anisotropic magnification; and (iii) per-particle defocus by CTF parameter fitting, including defocus fitting per-particle and astigmatism fitting per-micrograph. For the final refinement, the number of randomly selected particles used for each dataset was equalized to 6,488 particles for a fair comparison. The Rosenthal–Henderson *B*-factors were calculated as a quality metric using the bfactor_plot.py script (Zivanov *et al*., 2018) in RELION 5 (Table 2).

**Table 2.**
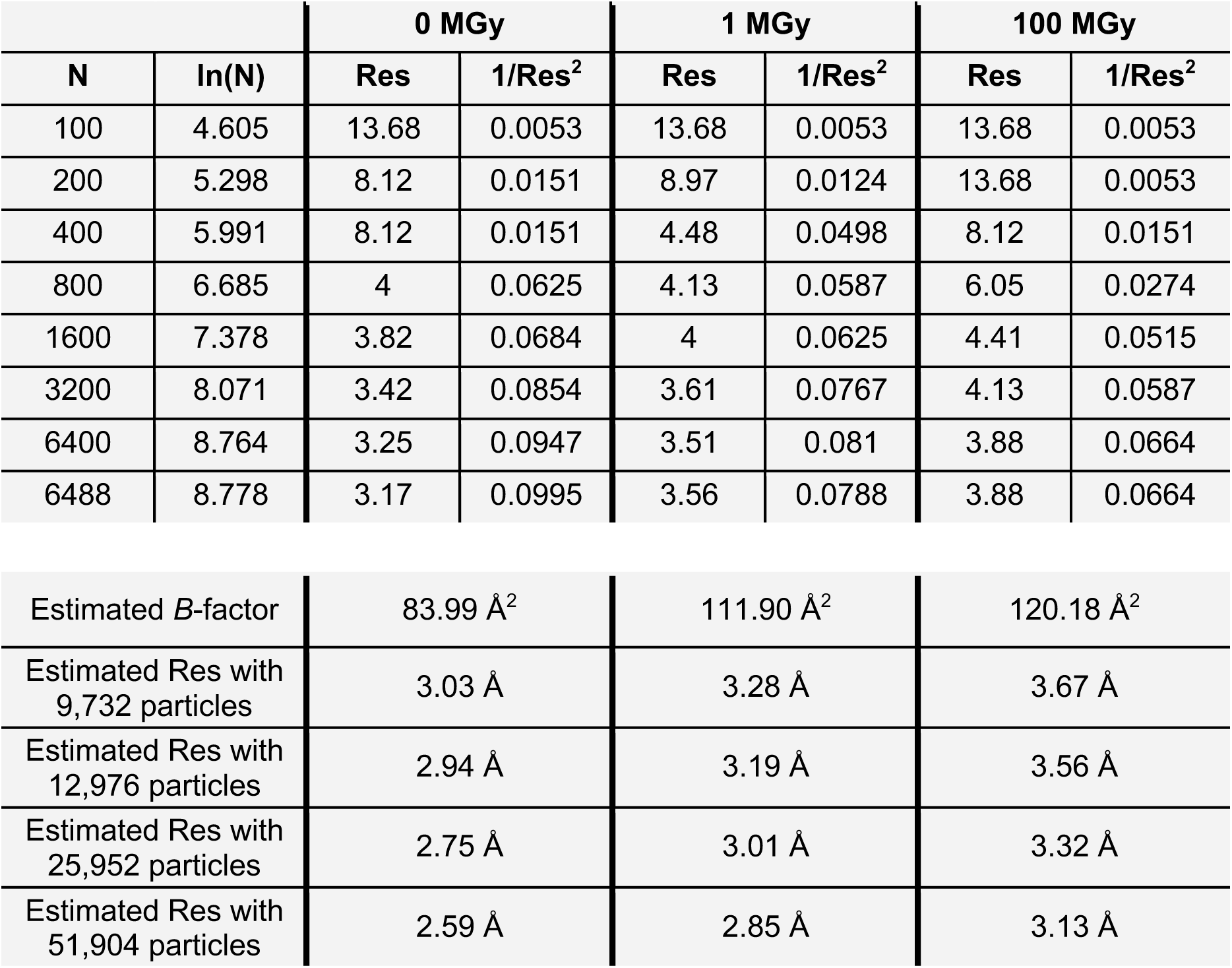
Rosenthal-Henderson *B*-factors (abbreviation Res = resolution).

|  |  | 0 MGy |  | 1 MGy |  | 100 MGy |  |
| --- | --- | --- | --- | --- | --- | --- | --- |
| N | ln(N) | Res | 1/Res<sup>2</sup> | Res | 1/Res<sup>2</sup> | Res | 1/Res<sup>2</sup> |
| 100 | 4.605 | 13.68 | 0.0053 | 13.68 | 0.0053 | 13.68 | 0.0053 |
| 200 | 5.298 | 8.12 | 0.0151 | 8.97 | 0.0124 | 13.68 | 0.0053 |
| 400 | 5.991 | 8.12 | 0.0151 | 4.48 | 0.0498 | 8.12 | 0.0151 |
| 800 | 6.685 | 4 | 0.0625 | 4.13 | 0.0587 | 6.05 | 0.0274 |
| 1600 | 7.378 | 3.82 | 0.0684 | 4 | 0.0625 | 4.41 | 0.0515 |
| 3200 | 8.071 | 3.42 | 0.0854 | 3.61 | 0.0767 | 4.13 | 0.0587 |
| 6400 | 8.764 | 3.25 | 0.0947 | 3.51 | 0.081 | 3.88 | 0.0664 |
| 6488 | 8.778 | 3.17 | 0.0995 | 3.56 | 0.0788 | 3.88 | 0.0664 |

|  |  |  |  |
| --- | --- | --- | --- |
| Estimated *B*-factor | 83.99 Å<sup>2</sup> | 111.90 Å<sup>2</sup> | 120.18 Å<sup>2</sup> |
| Estimated Res with 9,732 particles | 3.03 Å | 3.28 Å | 3.67 Å |
| Estimated Res with 12,976 particles | 2.94 Å | 3.19 Å | 3.56 Å |
| Estimated Res with 25,952 particles | 2.75 Å | 3.01 Å | 3.32 Å |
| Estimated Res with 51,904 particles | 2.59 Å | 2.85 Å | 3.13 Å |

## Results

### Exposing Apoferritin on EM grids to X-rays

Desalted apoferritin samples were applied to different grid types (see Materials and Methods), plunge frozen, and secured in AutoGrid Rings to enhance stability. These AutoGrid Rings were transferred on specialized 3D-printed holders for X-ray radiation exposure (Huang *et al*., 2020), stored under liquid nitrogen, and transported to the beamline ID30B at the European Synchrotron Radiation Facility (ESRF). Regions of the grids were exposed to X-ray radiation at a photon energy of 13.5 keV, absorbing doses of 1 and 100 MGy (see Methods for details on the dose calculation), which are within the range commonly used for X-ray nano-tomography. Following exposure, the samples were recovered under cryogenic temperature, shipped to the ScopeM facility at ETH Zurich, and analysed by cryo-EM and single particle analysis. Several issues arose during the experiment, including ice contamination, damage to the support film, damage to the grid bars (Figure 2), and, in some cases, the grids detaching from the holders. Of the two grid types tested, the more fragile Quantifoil Au200 R2/2 holey carbon EM finder grids did not survive handling, whereas the Quantifoil Cu200 R1.2/1.3 holey carbon EM grids withstood the mechanical stress. The specialized 3D-printed holders are not well suited for handling frozen cryo-EM grids (Figure 3A) under liquid nitrogen, as the plastic arms become rigid and brittle at cryogenic temperatures (Figure 3A). This creates difficulties when inserting the grid for two main reasons. First, if the holders are too tight, the grid cannot be inserted at all, requiring the holder to be warmed up before another attempt can be made. Second, if the holders are not sufficiently tight prior to cooling, the grid is inadequately secured and may fall out during handling, which could explain why, in some cases, grids became detached during shipment. Nevertheless, despite these challenges, one of the grids that had been exposed to 0, 1, and 100 MGy in different grid squares was still suitable for cryo-EM data collection.

**Figure 2.**
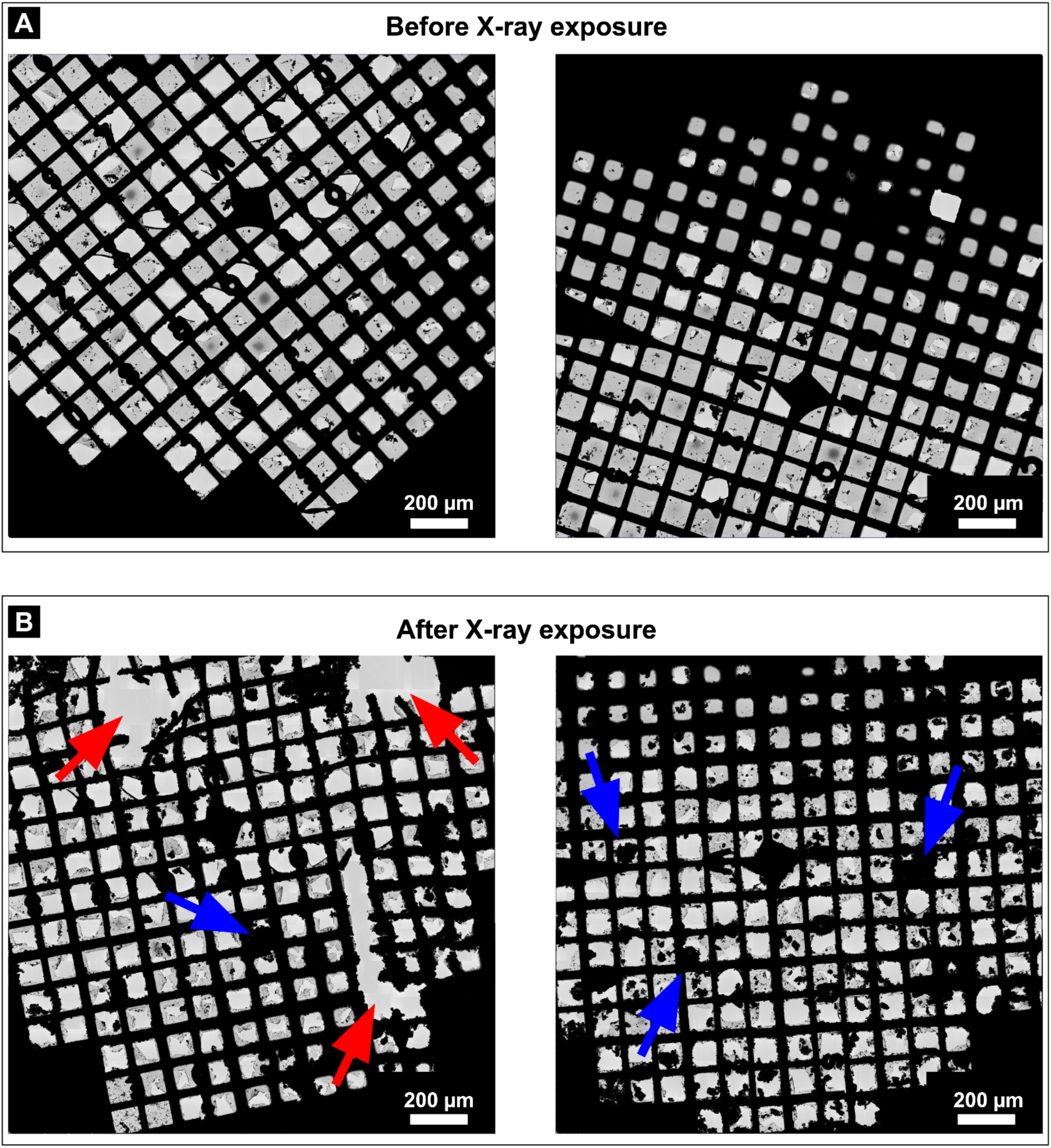
Cryo-TEM images of grids before and after X-ray irradiation. A-B. Representative of two grids before (A) and after (B) shipment to the synchrotron, exposure to X-rays, and shipment back in a dry shipper. The equivalent positions on the grids with either mechanical damage or ice contamination are indicated by red and blue arrows, respectively.

**Figure 3.**
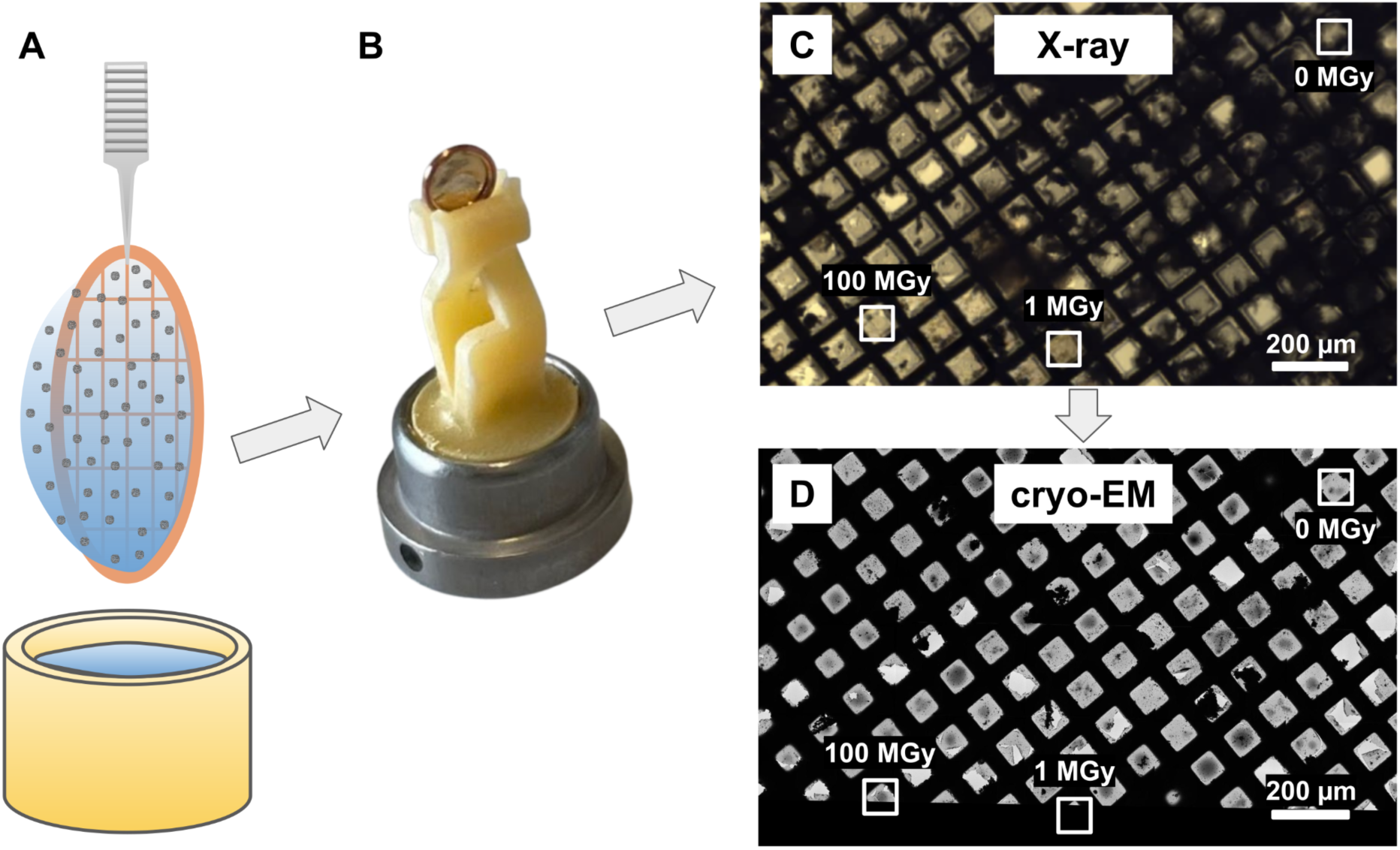
Correlation of synchrotron and cryo-EM imaging of frozen-hydrated apoferritin. (A) Apoferritin was plunge-frozen in liquid ethane. (B) The grid was clipped into AutoGrid rings and mounted onto custom 3D-printed holders. (C) At the beamline, the sample was transferred into the loading carrier, optically monitored with a camera and exposed to varying absorbed X-ray doses across selected grid squares. (D) The same grid squares were subsequently imaged by cryo-TEM, and data were recorded.

### Analysing apoferritin on exposed EM grids using cryo-TEM

Figure 3CD shows an atlas of the grid acquired by the cryo-EM, which was correlated with the image recorded at the X-ray crystallography beamline to locate the exposed areas. The processed grids displayed a high degree of ice contamination in comparison to the freshly prepared EM grids, but the grid remained in good condition relative to the other grids sent to the synchrotron (Figure 2). The relatively unbroken areas devoid of big ice clumps were selected for the final data acquisition. Even in the best regions of the grids, the micrographs displayed noticeable ice contamination (Figure S1A). Particles were automatically picked using a template and then subjected to multiple rounds of 2D and 3D classification to eliminate incorrectly picked, contaminated or poor-quality particles (Figure S1BC). This iterative cleaning ensured that only well-defined apoferritin particles were retained for further analysis. The selected particles were subsequently refined through CTF correction and polishing to improve data quality. To enable a fair comparison across conditions, the number of particles was normalized to 6,488 for each of the three data sets before final 3D refinement and post-processing. The resulting reconstructions demonstrated that, although higher radiation doses increasingly affected the samples, the high-resolution structural information was preserved in all cases, with each test data set yielding a robust reconstruction (Figure 4, Table 1).

**Figure 4.**
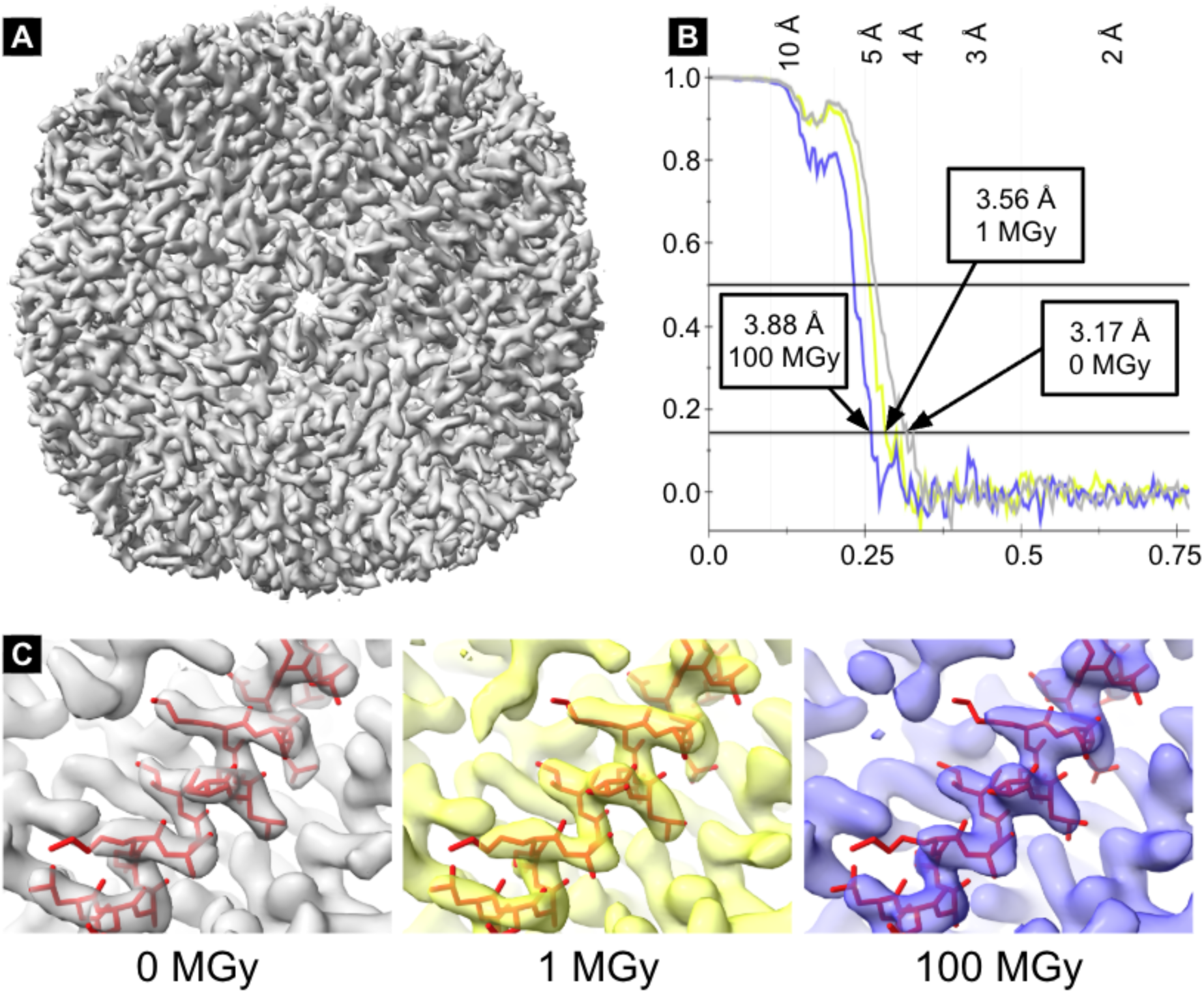
Apoferritin single-particle data analysis and final resolution estimation. (A) Complete electron density map of the apoferritin from 6,488 particles without X-ray exposure. (B) The Fourier shell correlation analysis of the three maps yielded resolutions of 3.17 Å for the dataset without X-ray exposure, and 3.56 Å and 3.88 Å for the apoferritin samples which had absorbed 1 MGy and 100 MGy X-ray doses, respectively. All three reconstructions were based on the same number of particles. These results indicate that higher X-ray exposure doses caused progressive protein damage, leading to a loss of resolution. (C) The apoferritin model (PDB 6rjh (Naydenova *et al*., 2019)) was fitted into all three maps to illustrate that each electron density map distinctly reveals the backbone and side chain densities, even though the side chain densities in the 100 MGy map appear noticeably weaker.

As expected, the particle selections from regions that had not been exposed to X-rays reached the highest resolution of 3.17 Å. Remarkably, the cryo-EM reconstructions of apoferritin that absorbed X-ray doses of 1 MGy and 100 MGy reached resolutions of 3.56 Å and 3.88 Å, respectively (Figure 4B). In line with this trend, the Rosenthal-Henderson *B*-factors also indicate diminished particle quality in the irradiated regions, rising from 83.99 Å^2^ at 0 MGy to 111.90 Å^2^ at 1 MGy and 120.18 Å^2^ at 100 MGy (Table 2). The level of detail preserved in the cryo-EM maps at this resolution would effectively allow macromolecular model building at near-atomic resolution, demonstrated in Figure 4C by the density quality that permits reliable fitting of the model (PDB 6rjh (Naydenova *et al*., 2019)). To ensure that the higher resolution was not simply an effect of ice thickness, micrographs were checked by eye and analysed with CTFFIND4 (Rohou & Grigorieff, 2015) for ice ring density (Figure S2). Both methods revealed more ice crystals for areas that had been exposed to X-rays. The pronounced increase in ice-related features in the exposed region, likely arising from both surface accumulation and local heating of the sample toward the devitrification temperature, probably accounts for the large number of particles excluded during 2D classification in the exposed area (82.3% and 83.0%), compared with the non-X-ray-exposed area (44.4%).

To check whether the resolution decrease was due to ice buildup, a dataset collected from a sample that had not been exposed to X-rays was processed without the first frame, corresponding to a pre-exposure of 1.5 e^-^/Å^2^ (≈ 5.55 MGy using a conversion factor of 3.7 (Dickerson et al., 2024; Groen et al., 2025)). The dataset only showed a slight increase in ice and a resolution drop of 0.25 Å to 3.42 Å (Figure S3). Taken together, these findings indicate that resolution changes result from the combined effects of ice accumulation and radiation damage. Importantly, they also demonstrate that even a combination of the highest absorbed X-ray dose of 100 MGy with a typical cryo-EM data collection workflow (accumulating a total dose of 60 e^-^/Å^2^ per movie stack) preserves the protein structure in this experimental format, i.e. purified protein in a thin film of vitreous ice, sufficiently for most structural biology applications at high resolution.

## Discussion

Multiscale correlative imaging of biological samples using X-rays and electrons has tremendous potential in light of the recent breakthroughs in both these techniques (Shafiei *et al*., 2025; Klein *et al*., 2021; Okolo *et al*., 2021; Lo *et al*., 2019). However, as both approaches are known to cause severe radiation damage to the objects of interest during imaging, a major question has been whether combining X-ray and cryo-EM analysis of intact frozen hydrated biological samples might destroy the atomic structures within the sample of interest, thereby rendering any meaningful high-resolution analysis largely impossible. Our results show that the absorption of a combination of a typical X-ray tomography radiation dose and a typical cryo-EM imaging dose preserves the high-resolution information in the sample, sufficient to reach near atomic 3D reconstructions for a model biological sample. When 100 MGy is converted to e^-^/Å^2^ at 300 keV using the factor of 3.7 (Dickerson *et al*., 2024; Groen *et al*., 2025), our findings are consistent with cryo-EM studies reporting that exposures of ∼19 e^-^/Å^2^ and ∼24 e^-^/Å^2^ at 300 keV yielded reconstructions of 3.35 Å (Grant & Grigorieff, 2015) and 3.94 Å (Allegretti *et al*., 2014), respectively, achieved with X-rays rather than electrons.

These results provide a strong foundation for developing a correlative imaging workflow combining cryo hard X-ray tomography and cryo-EM. Such a pipeline would allow multiscale imaging of thick biological specimens in a near-native frozen-hydrated state. In addition to validating this correlative imaging technique, our results, on one hand, reinforce the previous findings from X-ray crystallography regarding radiation-induced damage to protein crystal structures (Henderson, 1990; Gonzalez & Nave, 1994; Teng & Moffat, 2000; Owen *et al*., 2006), and on the other hand, provide a new perspective on studying X-ray radiation damage in the absence of a crystalline lattice as the major component. Remarkably, we show that a sample subjected to an intense X-ray dose of 100 MGy still yields a resolution of 3.88 Å when imaged via cryo-EM. This finding is highly significant as it suggests that structural integrity can be preserved even under harsh experimental conditions, broadening the range of feasible sample treatments in hybrid workflows.

The type of sample plays a key role: in this study, we demonstrated that thin samples used directly for cryo-EM lose resolution from both radiation damage and ice accumulation during X-ray exposure. By contrast, in workflows that combine hard X-ray nano-tomography with cryo-EM, thick samples are imaged at the synchrotron and then thinned to isolate regions of interest. Thinning removes the ice contamination introduced by X-rays, making these hybrid approaches especially effective.

However, there are limitations to our ability to interpret the results obtained with the thin model system here (i.e., apoferritin molecules suspended in a thin film of vitreous ice). It is conceivable that exposure of a much thicker frozen biological sample, such as a tissue biopsy, will result in the generation of more radicals due to X-ray interaction with the bulky specimen, leading to substantially more radiation damage than we have observed here. Careful experimentation will be needed to determine whether there are fundamental differences in the damage suffered by the thin and thick amorphous biological specimens.

Throughout the course of this research, we encountered two practical challenges while switching between X-ray microscopy and cryo-EM that warranted attention: (i) mechanical damage to the EM grid resulting from improper handling or exposure procedures, including incorrect grid orientation and autoloader miscalibration or misalignment; (ii) the presence of ice contamination arising during sample transfer. While these issues impacted the initial tests, they are unlikely to persist in the finalized protocol. To mitigate these risks, the refined workflow will use a larger biological sample embedded in a high-pressure frozen block mounted on a copper tube. This setup offers increased mechanical stability during imaging and transfer procedures. Following X-ray analysis, thin lamellae or cryo-sections will be prepared from the internal regions of the block - areas shielded from direct exposure in the X-ray microscope - thereby greatly reducing the risk of ice contamination and preserving sample quality for EM imaging. We anticipate that additional challenges will emerge, such as effective identification of the ROIs for lamella preparation via FIB or sectioning via CEMOVIS.

Together, these findings present a compelling case for a robust, integrated workflow bridging cryo hard X-ray microscopy and cryo-EM without compromising high-resolution data from biological samples. The demonstrated compatibility of the two modalities, along with practical solutions for minimizing damage and contamination, as well as a great potential for integrating additional imaging modalities, such as light microscopy, to facilitate targeting of the regions of interest for high-resolution cryo-EM imaging, positions this approach as a powerful tool for enhancing imaging workflows and future structural biology investigations. By leveraging X-ray tomography to map the frozen intact tissues and to identify biologically relevant regions, researchers may be able to apply cryo-ET or single particle analysis with near-atomic precision, thus expanding the capabilities of electron microscopy in studying complex biological systems.

## Acknowledgements

We acknowledge the support and assistance of the staff at beamline ID30B at ESRFwith particular thanks to Andrew McCarthy, and at the ScopeM facility at ETH Zurich, with special thanks to Miroslav Peterek and Bilal Qureshi. We also thank Spencer Bliven and Greta Assmann (PSI) for their support with high-performance computing.

## Data availability

The datasets have been deposited in the Electron Microscopy Data Bank (EMDB) under accession numbers 55343 (0 MGy), 55338 (1 MGy), and 55344 (100 MGy).

## Supporting information

**Figure S1.**
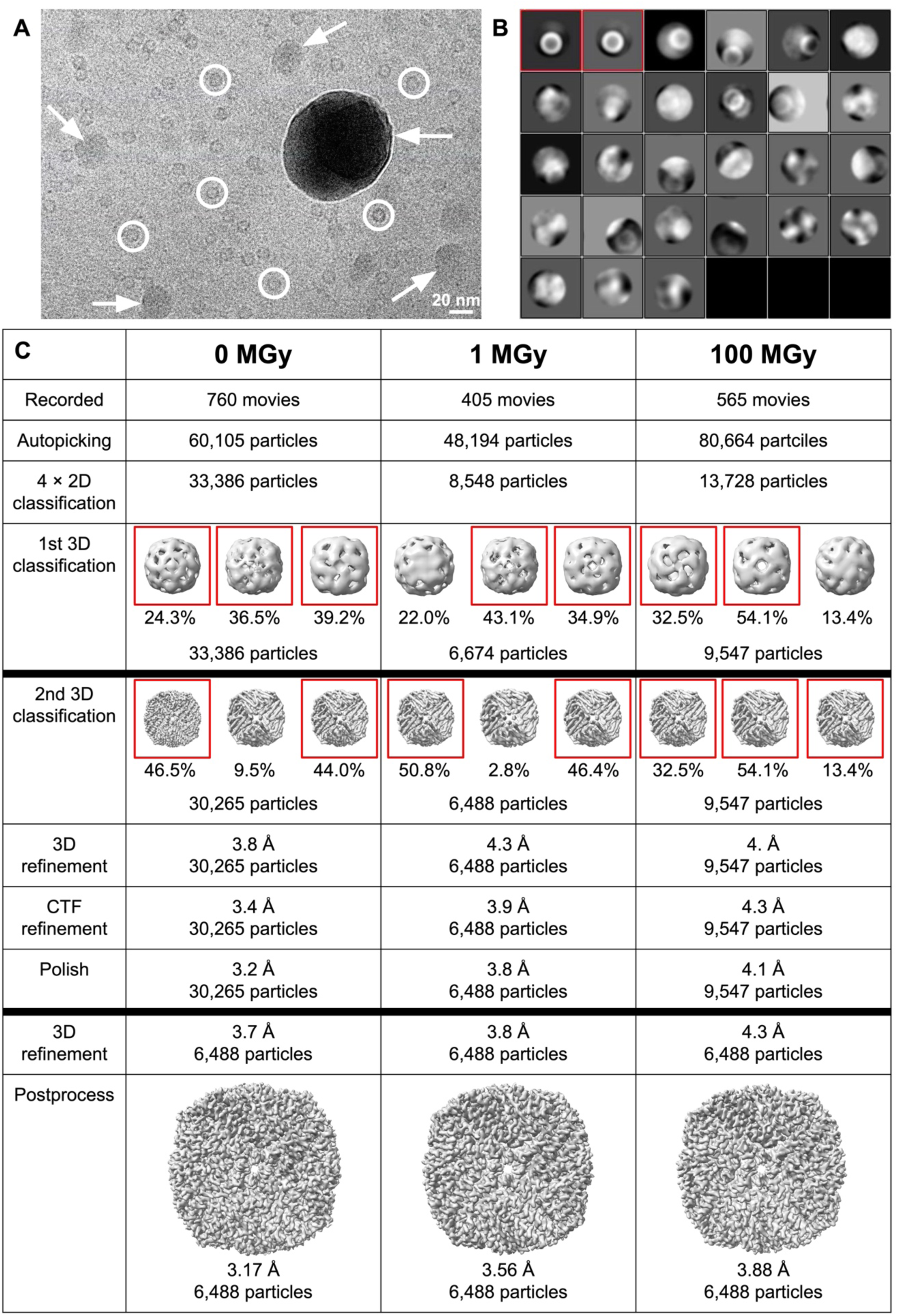
Processing pipeline of the three apoferritin data sets. (A) Representative micrograph showing apoferritin particles (circles) and regions of variable ice contamination (arrows). (B) Particles were template-picked and subjected to iterative 2D classification. Only classes with clear apoferritin features (red boxes) were retained. (C) Initial 2D and 3D classifications were performed with binned particles, followed by re-extraction at full resolution, final 3D classification, CTF refinement, and polishing. For comparison, each data set was equalized to 6,488 particles before final refinement and post-processing, yielding a resolution of 3.88 Å even after absorbing 100 MGy X-ray dose.

**Figure S2.**
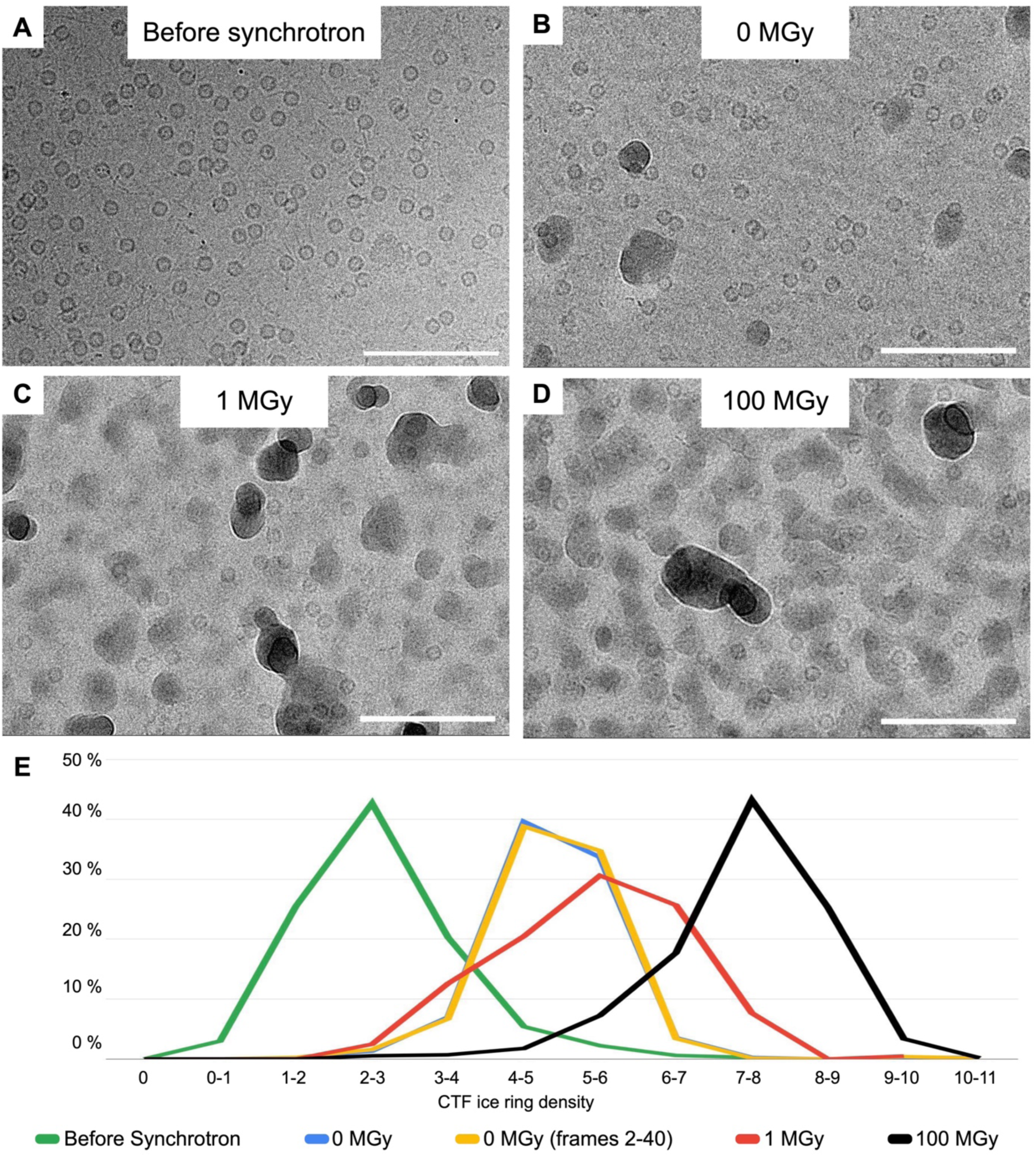
Ice contamination of samples. (A) Prior to synchrotron exposure, the sample displayed minimal crystalline ice. (B) After synchrotron handling, non-exposed regions showed increased ice, while areas that had absorbed X-ray doses of (C) 1 MGy or (D) 100 MGy exhibited even greater contamination (scale bars: 100 nm). (E) Quantification of CTF ice ring density using CTFFIND confirmed these observations, indicating that synchrotron handling promotes ice accumulation, which intensifies under X-ray irradiation.

**Figure S3.**
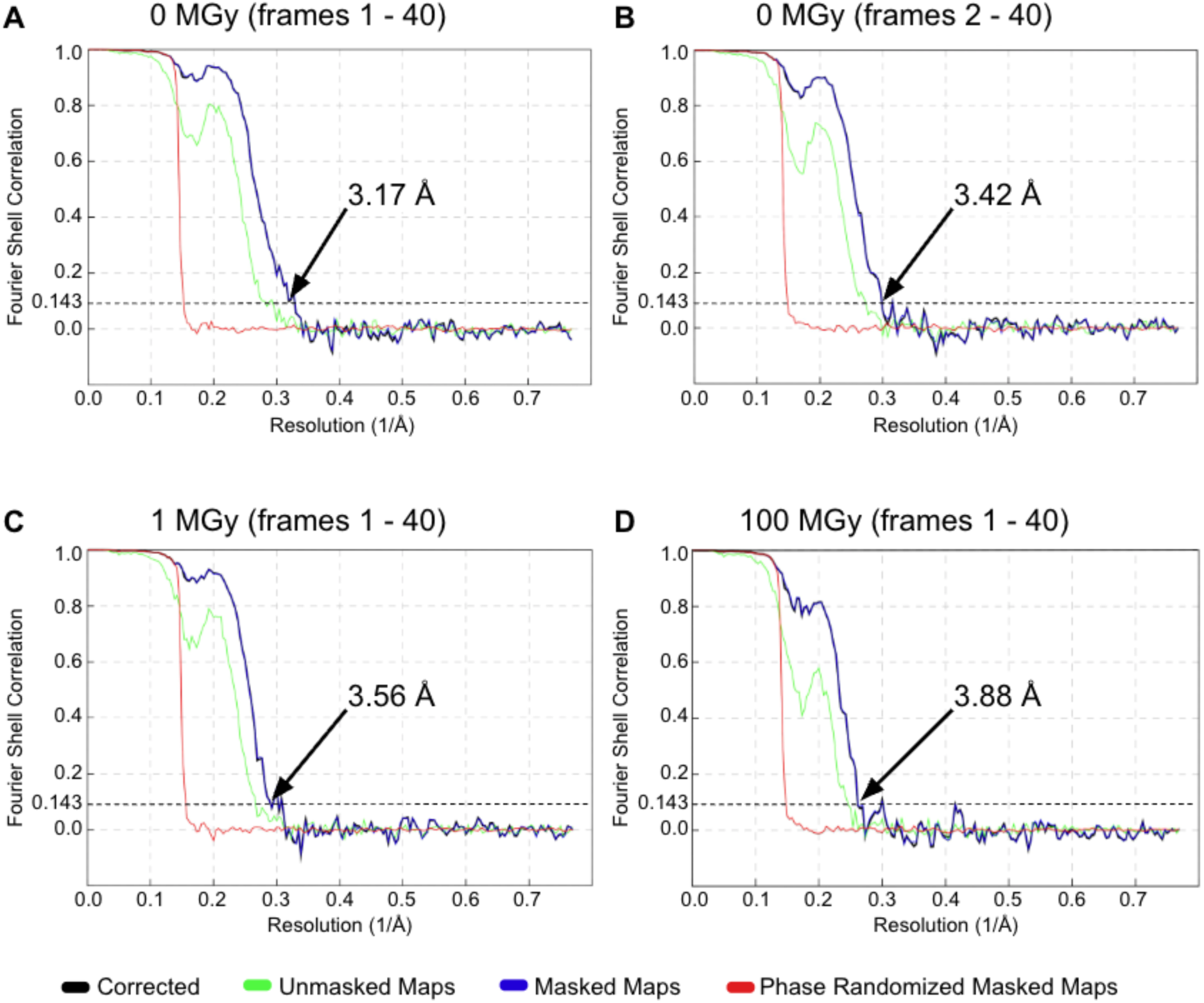
Fourier shell correlations. Fourier shell correlation (FSC) plots of apoferritin from a grid square not exposed to X-rays, using either (A) all frames or (B) excluding the first frame, yielded the highest resolution. In contrast, FSC plots from grid squares exposed to X-ray doses of (C) 1 MGy or (D) 100 MGy showed reduced resolution.

